# Single-cell RNA-Seq Resolves Cellular Heterogeneity and Transcriptional Dynamics in Spermatogonial Stem Cells Establishment

**DOI:** 10.1101/194696

**Authors:** Jinyue Liao, Shuk Han Ng, Jiajie Tu, Alfred Chun Shui Luk, Yan Qian, Jacqueline Fung, Nelson Leung Sang Tang, Bo Feng, Wai-Yee Chan, Pierre Fouchet, Tin-Lap Lee

## Abstract

The transition of gonocytes to spermatogonia and subsequent differentiation provide the foundation of spermatogenesis. However, systematic understanding on the cellular and molecular basis of this process is still limited, mainly impeded by the asynchrony in development and the lack of stage-specific markers. Using single-cell RNA sequencing on Oct4-GFP+/KIT- cells isolated from PND5.5 mice, we dissected the cellular heterogeneity and established molecular regulations. We demonstrated that gonocyte-spermatogonial transition was characterized by gene expression change related to apoptosis, cell cycle progression, and regulation of migration processes. Pseudotime analysis reconstructed developmental dynamics of the spermatogonial populations and unraveled sequential cellular and molecular transitions. We also identified CD87 as a neonatal stem cell marker which are potentially involved in the intial establishment of SSC pool. Lastly, we uncovered an unexpected subpopulation of spermatogonia primed to differentiation within the undifferentiated compartment, which is characterized by the lack of self-renewal genes and enhanced Oct4 expression and retinoic acid signaling response. Our study thus provides a novel understanding of cellular and molecular changes during spermatogonial establishment.

## INTRODUCTION

In mammals, male fertility is sustained by male germline stem cells (GSCs) known as spermatogonial stem cells (SSCs) that either self-renew or differentiate to progenitors to initiate spermatogenesis (Oatley and Brinster, 2008). The current model suggests that the initial pool of SSC is derived from the transition of postnatal gonocytes (Kluin and Derooij, 1981). Disturbances in cell fate decision during this process can lead to significant clinical consequences. In cryptorchidism, gonocyte transition into SSC is thought to be inhibited, which may result in infertility after puberty (Kamisawa et al., 2012). Defective SSC formation from gonocytes might be the cause of testicular germ cell tumors (TGCT) (Rajpert-de Meyts and Hoei-Hansen, 2007). SSC self-renewal and differentiation must also be sophistically balanced to maintain normal spermatogenesis and avoid tumorigenesis (Ishii et al., 2012).

The precursors of SSC pool arise from primordial germ cells (PGCs) during embryogenesis. The PGCs then differentiate to gonocytes, and quickly enter quiescent state after E13.5. Shortly after birth, gonocytes resume mitotic activity and begin migrating to the basal lamina of the testicular cords. Once they become resident at the basement membrane, the cells become spermatogonia including self-renewing SSCs (Culty, 2009; Manku and Culty, 2015; Saitou and Yamaji, 2012). A major challenge in deciphering the cellular and molecular mechanisms during this process is that the germ cells in neonatal testes are not developmentally synchronized, resulting in the coexistence of germ cells at different developmental stages (Nagano et al., 2000). It is impossible to visually distinguish the co-existing gonocytes and spermatogonia at various stages as they share similar morphology (Culty, 2009; Manku and Culty, 2015) and there is significant heterogeneity regarding marker expression among the germ cell populations (Hermann et al., 2015; Nakagawa et al., 2007; Suzuki et al., 2009; Yoshida et al., 2006b). This makes the population-based approach for molecular characterization infeasible, thereby hampering the pace of research in this field.

To resolve cellular heterogeneity and capture developmental dynamics, we have undertaken single-cell RNA-Seq to tackle this long-standing question through revealing biologically intermediate states of germ cell development from a single chronological time point (Li et al., 2016; Llorens-Bobadilla et al., 2015; Shin et al., 2015). We profiled 71 single-cell transcriptomes of germ cells from mouse testes on postnatal day (PND) 5.5, which constitute a mixture of gonocytes undergoing transition into spermatogonia and undifferentiated spermatogonia, including a subset of stem cells and committed progenitors.

This has allowed us to assign single cells to distinct cell states and reconstruct their developmental chronology. In doing so we delineated the molecular cascades governing gonocyte-spermatogonial transition (GST) and the balance of SSC self-renewal and differentiation. Our analysis further identified novel neonatal spermatogonial stem cell markers CD87. Lastly, we found that elevated Oct4 expression level correlated with higher Rarg expression and enhanced RA responsivieness, which signified the priming of undifferentiated spermatogonia differention.

## RESULTS

### Single-cell transcriptioanal profiling of germ cells from mouse testes at PND5.5

GST occurs between PND3 and PND6, during which gonocytes either give rise to the undifferentiated spermatogonia or directly into differentiating spermatogonia that enter the first wave of spermatogenesis (de Rooij, 1998; Yoshida et al., 2006a). However, the exact timing of the formation of the SSC pool has not been established. We aimed to characterize the developmental stage when germ cells consist of all the cell types: gonocytes, primitive spermatogonia, and differentiating spermatogonia. We chose PND5.5 as earlier time point will have only gonocyte and primitive spermatogonia while later without gonocytes. First, we set out to examine whether germ cells in PND5.5 testis would allow capture of the dynamic event of gonotyte-to-spermatogonia transition. Oct4-GFP transgenic mouse line is a great tool to study early germ cell development as it allows visualization of endogenous *Oct4* expression, which is restricted to undifferentiated germ cells at prepubertal stages (Ohbo et al., 2003). We found that majority of the Oct4-GFP+ cells were localized at the basement membrane, representing early spermatogonia including SSC. We also found cells resided within the center of testicular cords or positioned adjacent to the basement membrane with protrusions towards the periphery (Figure 1). These results confirmed the existence of transitioning gonocytes yet to finish the migratory phase at PND5.5. We then used several known markers to further define Oct4-GFP+ cells. Oct4-GFP+ cells exhibit significant overlap with PLZF, a marker for the undifferentiated state (Figure 1A, D), suggesting Oct4-GFP and PLZF expression are highly analogous and Oct4-GFP+ cells indeed label most, if not all, germ cells at this stage. Coexpression of Oct4-GFP with the important self-renewal factor GFRA1 was also observed in a subset of Oct4-GFP+ cells (Figure 1B, D). In addition, significant numbers of Oct4-GFP+ cells begin to express the differentiation marker KIT(Figure 1C). These results suggest that PND5.5 testis contains differentiating germ cells. Taken together, Oct4-GFP+ labelled germ cells at this single developmental time point should represent a continuum of differentiation from gonocytes to differentiating spermatogonia and single-cell transcriptome profiling of these cells would allow reconstruction of GST and subsequent differentiation *in vivo* (Figure S1A).

**Figure 1.**
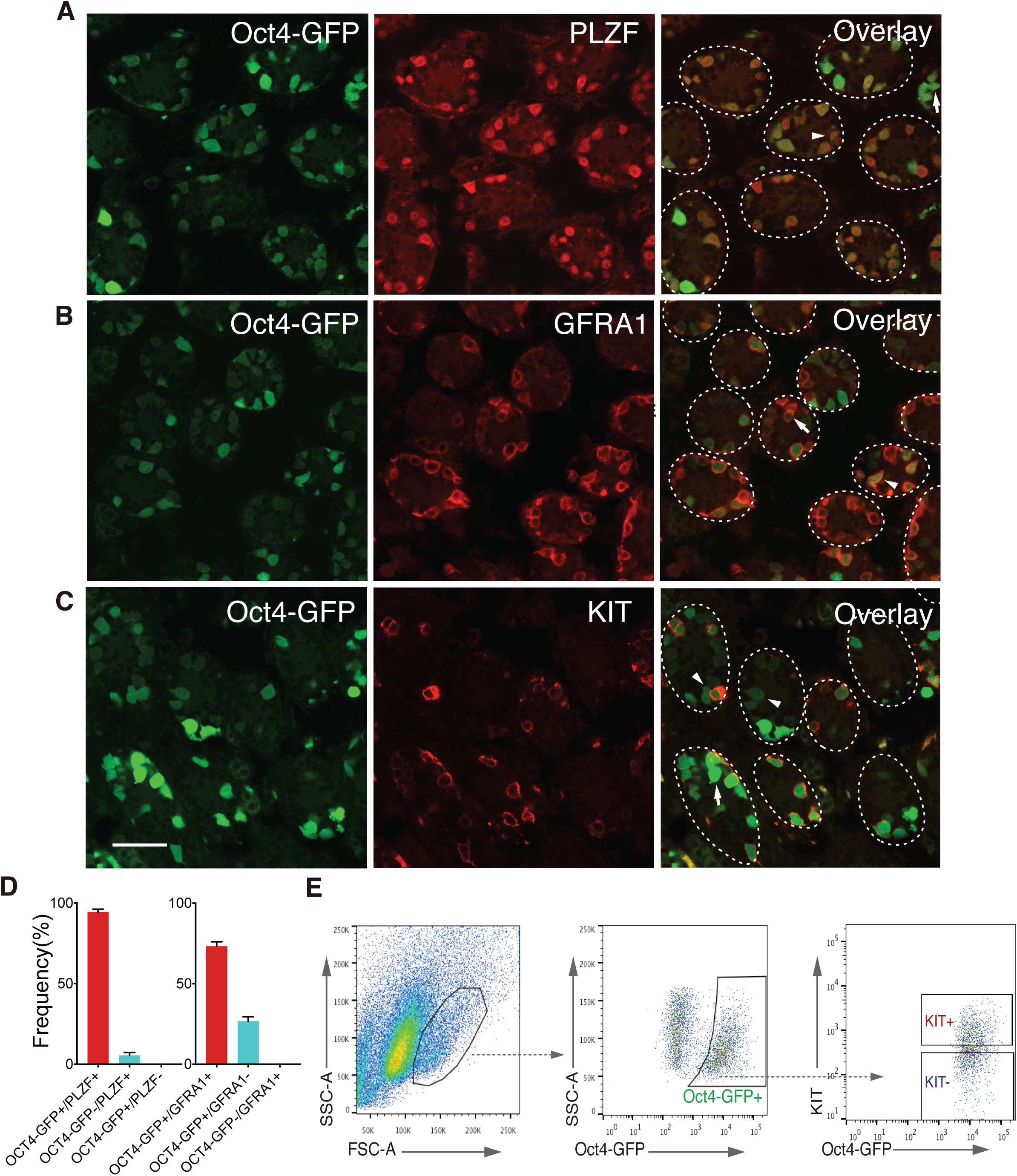
Characterization and isolation of Oct4-GFP+ germ cells from mouse testes at PND5.5. (A-C) Representative images of testicular sections from PND5.5 Oct4-GFP transgenic mice showing immunofluorescent staining of PLZF (A), GFRA1 (B), and KIT (C). Oct4-GFP expression was observed in cells that were co-stained with undifferentiated spermatogonial marker GFRA1 and PLZF (A and B). Oct4-GFP expressing cells can be classified as KIT- and KIT+ (C). Transitional gonocytes reside in the center of the testicular cord (arrow) or near the basement membrane with cytosolic projection towards the periphery (arrow head).Overlap of Oct4-GFP with PLZF or GFRA1 in testis section immunostaining result as shown in (A) or (B). Results are expressed as means ± SE of 4 testis sections. (E) Sorting strategy for isolation of undifferentiated (Oct4-GFP+/KIT-) at PND5.5. Scale bar = 50um.

To maximize the representation of the most undifferentiated germ cells, we isolated Oct4-GFP+/KIT- cells from PND5.5 testis using fluorescence-activated cell sorting (FACS) (Figure 1E, S1B-E). Single cells were then captured by Fluidigm C1 IFC system, which were subjected to automated RNA isolation, cDNA synthesis, library construction and RNA-sequencing (Figure S2A). 71 cells passed a number of stringent quality control assessments that ensured high-quality single-cell transcriptome data and were included in subsequent analyses (Figure S2B-F, Table S1). Although *Oct4* transcripts were absent in 19 of the 71 sequenced cells, likely because the expression was below the threshold of detection at this sequencing depth, *Plzf* transcripts were abundantly expressed in all cells, indicating their germ cell nature (Figure S3). Furthermore, a significant portion of germ cell-related genes such as *Bcl6b*, *Egr2*, *Sohlh1*, and *Csf3r* showed substantial variability, while housekeeping genes (*Actb*, *Gapdh*, *Rpl7*, and *Rps2*) showed uniform expression across single cells, illustrating the significant gene expression heterogeneity in early male germ cell populations as previous reported (Hermann et al., 2015) (Figure S3).

### Single-cell RNA-Seq analysis recapitulates germ cell subpopulations and cellular dynamics

We next sought to interrogate the heterogeneous cellular composition. Genes with the highest loadings in principal component analysis (PCA) were analyzed by unsupervised hierarchical clustering, which identified four different subpopulations (Figure 2A, B), including 2 smaller clusters (I and IV) and 2 larger clusters (II and III). Aforementioned characterization implies that majority of cells captured should be spermatogonia at PND5.5 and may be linked to Cluster II and III. Visual inspection of the heatmap suggests that distinctions among Cluster II and III appeared to predominantly reflect a gradient of expression in mRNA levels, suggesting the two subsets of spermatogonia could represent temporally distinct populations. Thus, Cluster II cells with higher expression of genes associated with SSC self-renewal, such as *Etv5* and *Ccnd2* (Figure 2C), are likely to be the primitive stem cells. In contrast, Cluster III cells, which display higher expression of differentiation regulator *Sohlh1*, are the immediate progenitors (Figure 2C). The expression level of *bona fide* adult stem cell markers in Cluster II and III also reflected their cellular identities (Abid et al., 2014; Aloisio et al., 2014; Oatley et al., 2011). The majority of cells with high *Id4* expression fell into Cluster II (15 out of 23 cells, log2(TPM)>10)), consistent with the recent observation that cells with high level of *Id4* represent true SSCs ((Helsel et al., 2017)). *Erbb3* and *Pax7* were also preferentially expressed in this primitive stem cell pool (Figure 2C). We further defined Cluster IV cells as differentiation-primed spermatogonia, based on their close proximity in the PCA space to the Cluster III progenitors and the absence of *Gfra1* (Figuer 2B and C), which are likely to be primed to differentiation and lose self-renewal potential (Meng et al., 2000). The last cell subset (Cluster I) identified in the present experiment corresponds to the gonocyte, based on the PC2 distance between them and the spermatogonia (Cluster II, III and IV) and the fact that existence of gonocyte population should be reflected in single-cell transcriptomes of the captured cells (Figure 2B). It is also notable that *CD81*, which is reported to mediate migratory gonocyte movement (Tres and Kierszenbaum, 2005), showed higher expression in Cluster I (Mean TPM: Cluster I=62, cluster II=18, Cluster III=22, and Cluster IV=10) (Figure 2C). A single-cell characterization of germ cells in the neonatal testis was recently published using a different reporter ID4-GFP (Song et al., 2016). This provides a unique opportunity to explore the relationship among germ cells at different developmental stages. When we projected cells from our study onto this reference dataset, we observed that Cluster I cells bear a property intermediate between that of P3 gonocytes and P5.5 spermatogonia (Cluster II), while Cluster III and IV cells more closely resemble the P7 spermatogonia, affirming the gonocyte nature of Cluster I cells (Figure 2D). We further reconstructed the transitional dynamics and ontogeny relationship of the cell populations by Monocle (Trapnell et al., 2014b). The result supported our proposed assignments, where the trajectory path began with transitional gonocytes (Cluster I), passing through undifferentiated spermatogonia (Cluster II and III) and ending at differentiation-primed spermatogonia (Cluster IV) (Figure2E, left). Monocle also revealed three major development stages that might correspond to GST (State 1), SSS self-renewal (State 2), and differentiation commitment (State 3) (Figure 2E, right).

**Figure 2.**
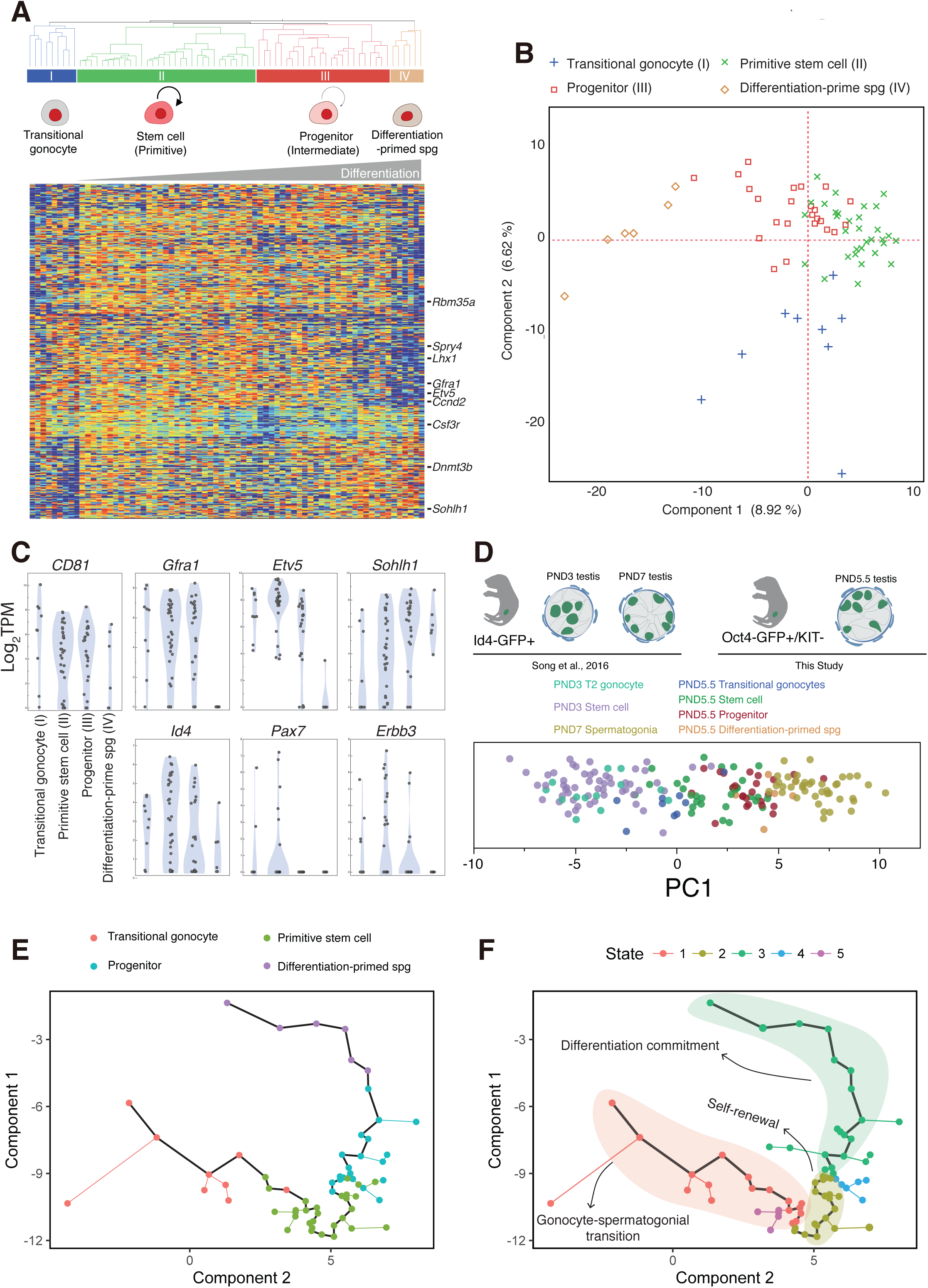
Single-cell transcriptom analysis reveals subpopulations and cellular transitions. (A) Hierarchical clustering analysis of 71 single-cell transcriptomes by Ward’s method uncovered 4 subpopulations. Germ cell subpopulation identities are resolved by known marker genes as shown in (C). (B) Principal component analysis of 71 single-cell RNA-seq transcriptomes. Each point represents a single cell and labelled according to clustering result as shown in (A). Data were plotted along the first and second principal components. (C) Violin plots illustrating the distribution of representative genes for subpopulations (top) as well as adult A_s_ marker genes (bottom). (D) Comparison of FACS sorting schemes implemented in our study and in the Song et al. study was shown at the top. PCA on all ID4-GFP+ transcriptome from Song et al. study using the expression [log2(TPM+1)] of the top500 PCA genes in our study was shown at the bottom. 71 single-cell transcriptomes in our study are projected onto the resulting principal component space. Cells colored by developmental identity indicated at the top. (E) Reconstructing continuous trajectory and assigning pseudotime for germ cell development using Monocle. Coloring of cells based on the group assignment from hierarchical clustering reveals stepwise development from gonocytes (Cluster I) to undifferentiated spermatogonia (Cluster II and to differentiation-primed spermatogonia (Cluster IV) (left). Developmental states revealed by Monocle (right).

We further validated our findings in a second set of PND 5.5 germ cells by single-cell RT-qPCR. We sorted an additional 96 cells and quantitatively measured expression of panel of genes that are preferentially expressed in each subpopulation using Biomark microfluidic quantitative PCR platform (Figure S4A-C, Table S2). Our analysis using these data showed that the pattern of PCA and hierarchical clustering followed the same structure as observed in the RNA-Seq experiment (Figure S4D and S4E). Similarly, the gene expression generally recapitulated patterns we identified in the initial RNA-Seq experiment (Figure S4F). These results demonstrated that the cell populations and their relative relationships revealed in the scRNA-Seq analysis are not due to biological or technical artifacts.

### Transcriptional regulation of gonocyte-spermatogonial transition

The hallmarks of perinatal gonocytes are their relocation from the center to the periphery of seminiferous cords and their cell cycle reentry (Nagano et al., 2000). To directly link Cluster I subpopulation identified in our data to gonocyte, we first assessed expression of migration-related genes within subpopulations. We focused on the cells resided in gonocyte-spermatogonial transition state as revealed by Monocle (State 1) (Figure 2E) and compared gene expression between the first half (9 cells from Cluster I) and the second half (9 cells from Cluster II) in the trajectory. Top-ranked differential expressed genes identified were postulated to regulate cell migration (Figure 3A). For example, this analysis revealed *CD81* as one of the top genes. Cluster I also expressed higher *Crk*, an adapter protein of PDGFR, which might be involved in PDGF-mediated gonocyte survival and migration (Basciani et al., 2008). Moreover, among genes up-regulated in the gonocytes, the top GO categories enriched was “Integrin”, which contained genes *Nf2*, *Itgb1*, *Utrn*, *Mfge8*, and *P4hb*. *Itgb1* (β1-integrin) has been implicated as an essential adhesion receptor required for both gonocytes migration and SSC homing (Kanatsu-Shinohara et al., 2008; Tres and Kierszenbaum, 2005). Interestingly, top-ranked genes enriched in migratory gonocytes included *Trpm7*, indicating its possible role in gonocyte migration through calcium-mediated cell movement (Chen et al., 2010).

**Figure 3.**
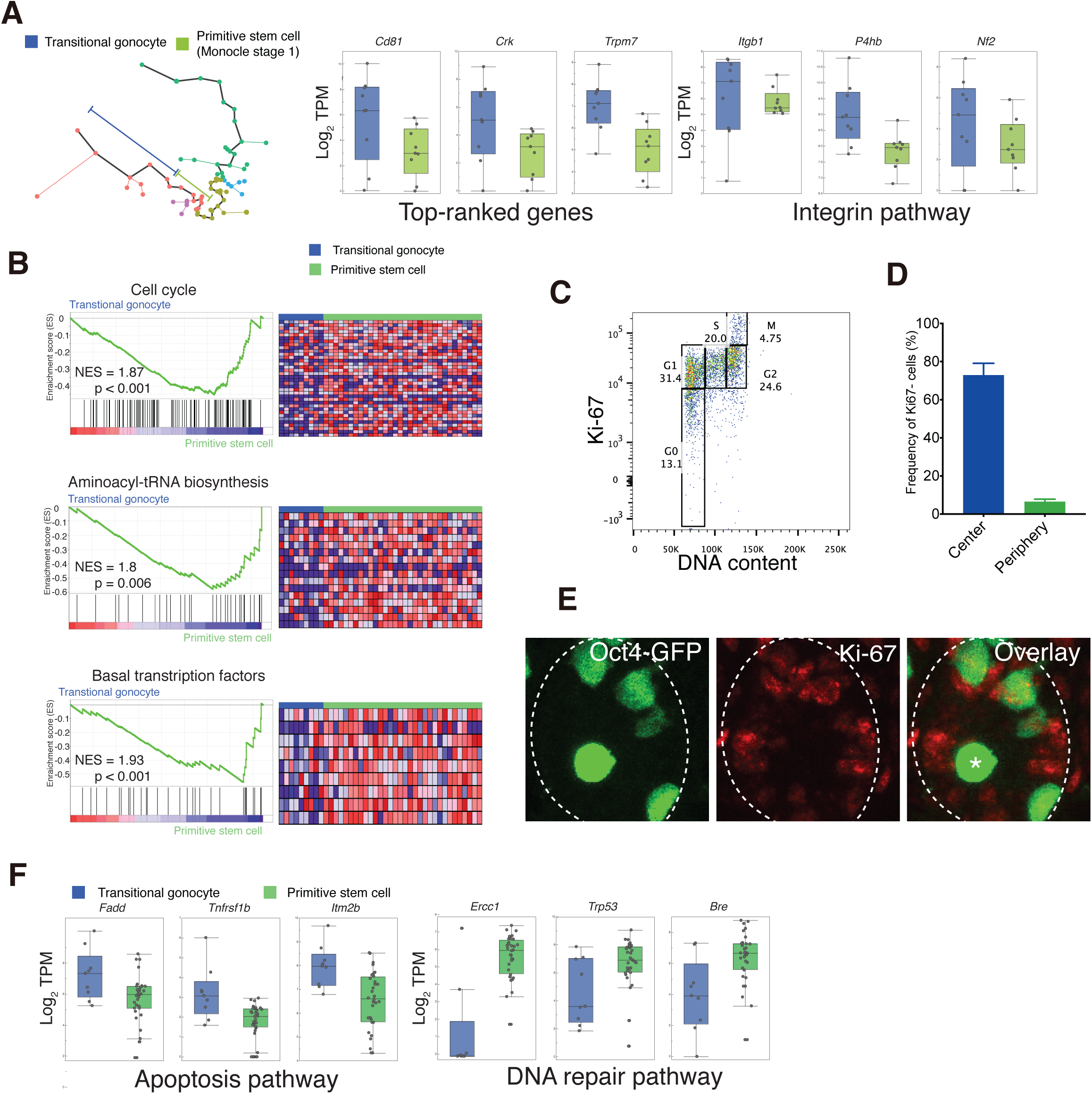
Characterization of gene expression and signaling pathways in gonocyte-spermatogonial transition. (A) Box plots for representative genes identified as differentially expressed (FDR<0.05) between cells in the first half and second half of stage 1 as indicated in Figure 2E, right. (B) Signaling pathways enriched in primitive stem cell as compared to transitional gonocytes by GSEA/KEGG analysis. Left, GSEA enrichment plot of KEGG signaling pathways. Right, heat maps of genes enriched in representative signaling pathways. The values indicated on individual plots are the Normalised Enrichment Score (NES) and p-value of the enrichment (permutation analysis). (C) Exemplary FACS plots of Ki-67 and Hoechst (DNA content) of Oct4-GFP+/KIT- cells to assess resting and cycling cells. (D) The percentage of Ki-67 negative cells in the cells located at the tubular basement membrane and at the center of the seminiferous tubules, showing a significantly higher percentage of Ki-67-cells in the centrally located population. (E) Cross-sectional images of Ki-67 (red) immunostaining of testis from Oct4-GFP mice. The outlines of seminiferous tubules are indicated by dashed lines. The asterisk denotes the cell resides at the center of the tubules and lacks Ki-67 expression. (F) Box plots for apoptosis-related genes identified as differentially expressed between transitional gonocytes and stem cell spermatogonia.

To further confirm the gonocyte nature of Cluster I, we perform Gene Set Enrichment Analysis (GSEA) and found that gene sets associated with cell cycle and protein synthesis (aminoacyl-tRNA biosynthesis) related pathways were underrepresented in the gonocyte population, implying a resting phase of these gonocytes prior to their transition to spermatogonia (Figure 3B). Interestingly, gonocytes were shown to express lower levels of basal transcriptional factors (TFIIB) like *Taf4b*, which encodes a chromatin-associated factor and has been demonstrated to regulate early germ stem cell specification (Falender et al., 2005). To experimentally validate the existence of this population of “cell cycle-low” cells, we examined the proliferation marker Ki-67 in Oct4-GFP+/KIT- cells. In line with the presence of 12.6% (9 in 71 cells) cells (Cluster I) within the cells captured, we observed a similar fraction of Oct4-GFP+/KIT- cells that were negative for the Ki-67 expression (Figure 3C). Immunostaining of the testis revealed that the centrally located cells contain greater proportion of Ki-67-negative cells than peripherally located cells (Figure 3D and 3E). GO analysis identified significant up-regulation of genes associated with death receptor pathway in gonocytes, which contained *Fadd*, *Tnfrfs1b*, *Icam1*, *Madd*, *Stx4a*, and *Skil*. We also found elevated expression of genes involved in intrinsic apoptosis signaling (*Itm2b*) (Figure 3F) (Fleischer et al., 2002). Genes associated with DNA damage/repair pathway were shown to be repressed in gonocytes, including *Ercc1*, *Trp53*, and *Bre* (Figure 3F). These results are in line with the notion that apoptosis plays an integral role in gonocyte cell fate determination (Wang et al., 1998).

Collectively, our analyses revealed transcriptional signatures that might be involved in cell fate decision and migration of germ cells during GST.

### Characterization of stem cell and progenitor spermatogonia by pseudotime ordering and network analysis

To investigate the underlying molecular features of the progression from the early, stem-like stage to the progenitor stage of spermatogonia, we analyzed differentially expressed genes between Cluster II and Cluster III. We identified 511 genes differentially expressed, which included *Etv5* and *Pax7* (Table S3). GO analysis revealed significant enrichment for signatures associated with integrin complex, Rap1 signaling and lipid metabolism in primitive stem cell pool (Figure S5A), whereas TGF signaling (e.g. Smad3 and Creb1) overrepresented in progenitor pool (Figure S5B). GSEA analysis further revealed that JAK/STAT signaling, cell adhesion molecules (CAMs), neuroactive ligand-receptor interaction, and RIG-I-like receptor signaling pathway were upregulated in stem cell cluster, while ribosome biogenesis pathway upregulated in progenitor subset (Figure 4A and S6A). To show coordination on gene expression during spermatogonia differentiation at PND5.5, we applied Monocle to position genes with similar expression trends along differentiation. We revealed three gene regulatory events following the self-renewal phase during spermatogonia differentiation (Figure 4B). During the initiation phase, known self-renewal genes such as *Ccnd2* and *Etv5* as well as signaling pathways, including MAPK/Erk1 and VEGF were repressed, as reported previously (Figure 4C) (Caires et al., 2009; He et al., 2008). Concurrently, a series of genes highly enriched for cell cycle and cell division was activated and then repressed before commitment of differentiation, suggesting an expansion phase precedes terminal differentiation of spermatogonia. The cells passed through an intermediate phase and transiently expressed an elevated level of *Nanos3* and genes related to DNA repairing pathway (*Bre* and *Trp53*) (Figure 4C). Previous work has indicated that *Nanos3* preferentially expresses in A_al_ spermatogonia that are committed to differentiation in adult testis (Suzuki et al., 2009). The commitment phase featured complete repression of SSC self-renewal genes and elevation of differentiation genes, such as *Sohlh1*, a point we will return to below. The interaction between cell cycle and differentiation prompted us to dissect the cell cycle stages by PCA-based approach (Buettner et al., 2015). We found that the majority of cells (70%, 14 cells out of 20) in G2/M phase were from Cluster II, from which we inferred that more cells in G2/M were sampled from primitive stem cell pool (Figure S6B). This implied that stem cells tend to have a relatively shorter G1 phase and proliferate faster, which is reminiscent of embryonic and adult stem cells (He et al., 2009; Lange et al., 2009).

**Figure 4.**
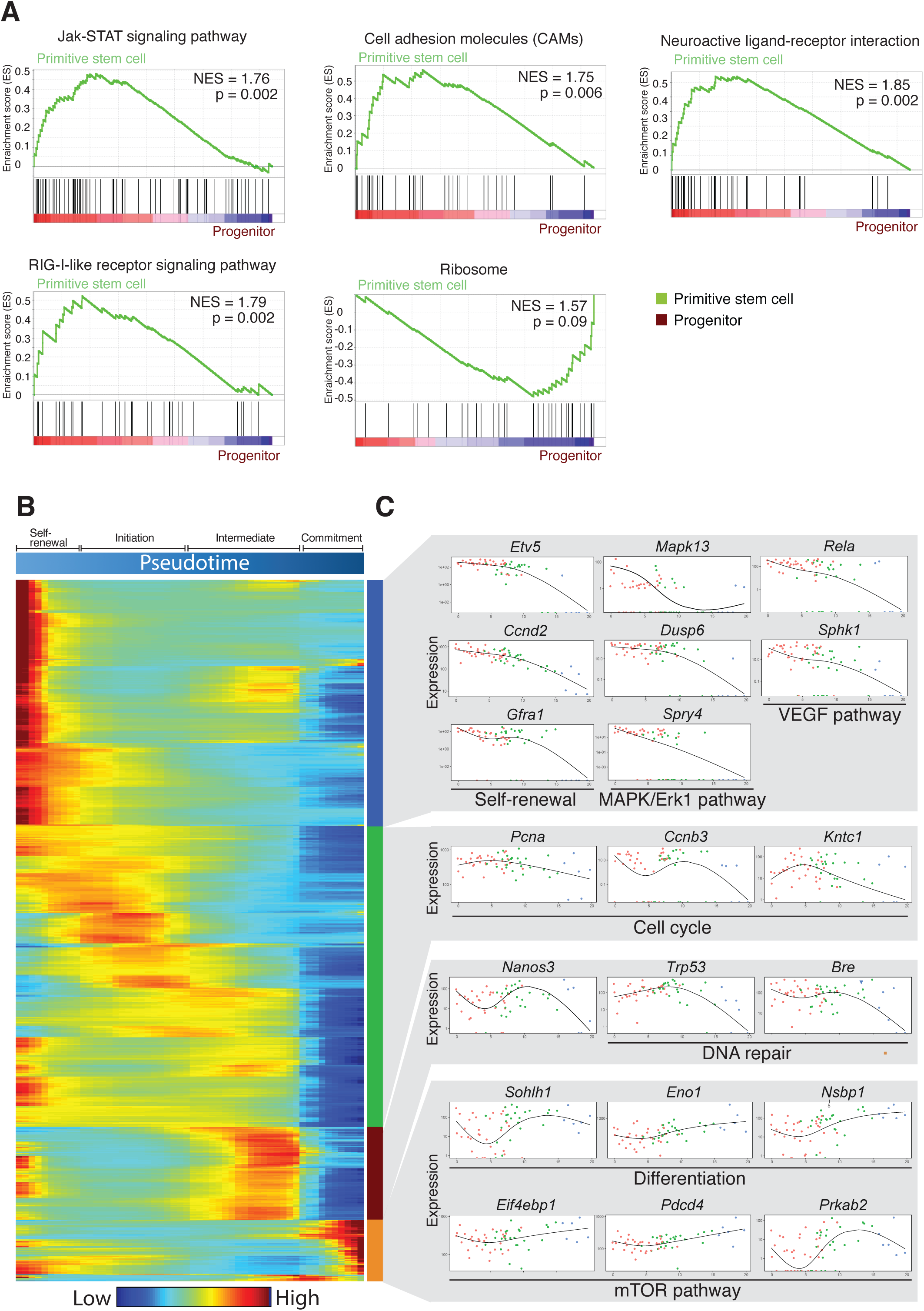
Molecular cascades regulating spermatogonia self-renewal and differentiation. (A) Signaling pathways enriched in stem cell spermatogonia as compared to progenitors by GSEA/KEGG analysis. The values indicated on GSEA enrichment plots are the Normalised Enrichment Score (NES) and p-value of the enrichment. (B) Hierarchical clustering of genes that significantly vary over pseudotime in spermatogonia differentiation process using expression kinetic curves calculated by Monocle. The bar on the right shows 4 distinct kinetic trends recovered. (C) Pseudotime profiles of representative genes showing different kinetics of expression.

To identify novel regulatory networks associated with SSC self-renewal and differentiation, we applied Weighted Gene Co-expression Network Analysis (WGCNA) algorithm on transcriptomes of spermatogonia (Langfelder and Horvath, 2008). In search for modules that correlate with our differentiation model, we identified the pink module that enriched in primitive stem cells and decreased across pseudotime (Figure S6C, left). GO terms of this module are associated with cell proliferation, regulation of developmental process and neuron projection. This unbiased approach highlighted key regulators as central to this network, including *Etv5*, *Ccnd2* and *Gfra1* as well as the presence of GDNF signaling regulation (19 out of 107 genes are GDNF-regulated), which confirmed the current knowledge of SSC self-renewal (He et al., 2009). It would be interesting to characterize functional significance of other highly connected hub genes such as *Tcl1*, *Spry4*, *Nefm*, *Dusp6*, *Tmem59l* and *Impact* (Figure S6C, left). The second module was enriched for genes involved in mitotic cell cycle and RNA binding. It was mostly repressed in differentiation-primed cells, suggesting a general reduction in proliferation rate upon differentiation commitment (Figure S6C, right). Hub-gene-network analysis indicated *Tex13* and *Rbm35a* are the most highly connected genes, both of which might exert potential functional roles in spermatogonia differentiation (Fagoonee et al., 2013; Kwon et al., 2014).

### CD87 serves as a neonatal spermatogononial stem cell marker

We next aimed to identify putative markers or regulators that may be specific to the earlier, potentially self-renewing spermatogonia (Cluster II) (Figure S7A). Surface marker *Tspan8* was also shown to correlate with stem cell activity in neonatal testis, which further validated our method (Mutoji et al., 2016). We found *CD87* particularly appealing as it has been implicated in hematopoietic cell migration and homing, and served as a marker for a subset of hematopoietic stem/progenitor cells (HSPCs) (Tjwa et al., 2009). We tested whether Cluster II-enriched *CD87* mRNAs truly reflected protein expression. Immunostaining signals of CD87 showed that they are more restrictedly expressed when compared to Oct4-GFP signal, which therefore establishes novel cellular heterogeneity within undifferentiated spermatogonia (Figure 5A). In line with development model we reconstructed, FACS-sorted cells positive for CD87 showed higher expression of *Etv5* and *Ccnd2*, and reduced expression of *Sohlh1* and *Ngn3* (Figure 5B-C, S7B). The foregoing analysis predicted that primitive stem cells tend to have a relatively shorter G1 phase and proliferate faster. Therefore, we investigated whether CD87+ cells, as a proxy to primitive stem cells, differed from CD87-cells in regard to cell cycle activity. FACS analysis confirmed a higher frequency of cells in G2/M phase in CD87+ versus CD87-cells, likely reflecting a faster proliferation rate (Figure 5F). Indeed, we observed significantly higher growth rate for germ cell (GS) culture derived from CD87+ cells (Figure 5G). To gain more insight of CD87’s function in germ cells, we examined its surface expression at postnatal stages by FACS. All Oct4-GFP+ cells expressed CD87 at early stages and the proportion gradually decreased from PND4.5 to PND7.5 and became nearly undetectable by PND14 (Figure 5D). We next explored whether CD87 participated in germ cell migration. We performed matrix adhesion assays using GS cultured on matrigel and examine the formation of the cellular protrusion, which has been described as an essential step for cell migration (Ladoux and Nicolas, 2012). Interestingly, CD87+ cells display cellular protrusion more frequently than CD87-cells, suggesting a greater migration potential (Figure 5E). Collectively, these results indicated that CD87 participated in spermatogenesis shortly after birth and played a specialized role in facilitating the establishment of the first SSC pool.

**Figure 5.**
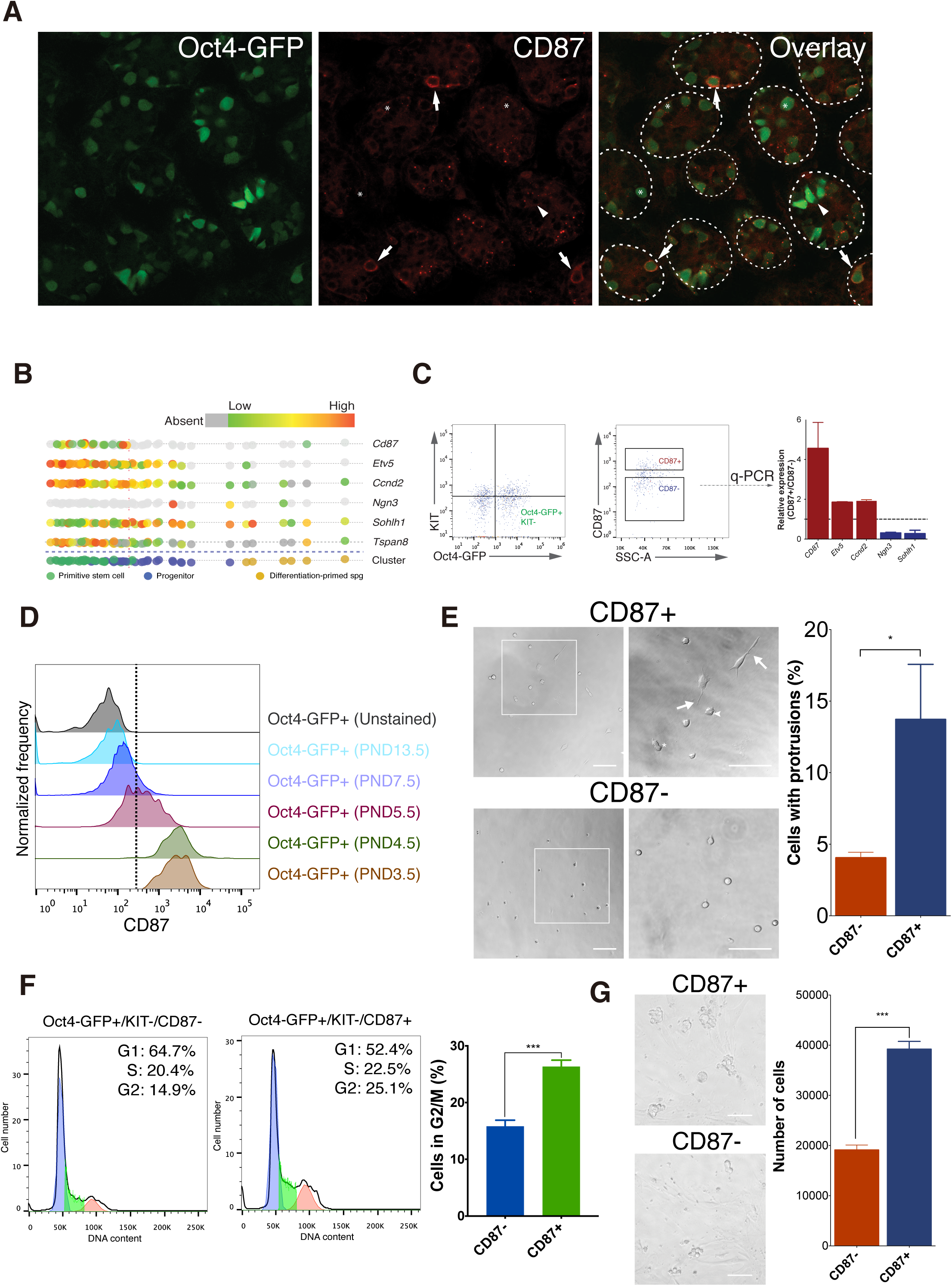
CD87 is a novel surface marker for primitive stem cells. (A) Confocal image of testis section from mouse at PND5.5 immunostained for CD87. Arrows indicate cells with strong CD87 expression. The asterisks and arrowheads denote spermatogonia and transitional gonocytes lacking CD87 expression, respectively. (B) The computationally reconstructed temporal gene expression pattern of genes tested. Each point corresponds to one cell and the color on each cell corresponds the scaled expression value of the respective gene. (C) Validation of gene expression patterns in CD87+ and CD87-subpopulations by FACS and qRT-PCR. (D) The CD87 expression is dynamic throughout the spermatogenesis initiation. Oct4-GFP cells were isolated at 3.5–14 days after birth and stained with APC-conjugated anti-CD87 antibody. The histogram plots show the expression of surface CD87 at different time points. The dashed line represents the cutoff for the determination of CD87+ v.s. CD87-based on staining with isotype controls. (E) Brightfield images showing that CD87+ cells display more spreading and lamellipodia formation on Matrigel. Quantification of cell protrusion comparing CD87+ and CD87-cells is shown on the right. (F) The representative result of cell cycle distribution analysis comparing the CD87+ and CD87-cells. Distribution of cells in G1, S and G2/M phases of the cell cycle as determined using FlowJo software. The percentages of cell populations of different cell cycle phases are shown in the upper-right corner. Quantification on the right showing a significantly higher percentage of G2/M cells in the CD87+ population. (G) The appearance of GS cell cultured derived from freshly isolated CD87+ and CD87-cells. 10000 FACS-sorted cells were seeded in each well. The representative brightfield images taken 3 days after seeding were shown on the left. The quantification shows significantly higher proliferation rate of CD87+ cells.

### Gene expression changes during early spermatogonia differentiation

To identify the molecular regulation of spermatogonia differentiation, we compared gene expression between “differentiation-primed” (*Gfra1*-/*Kit*-, Cluster IV) population and its undifferentiated counterpart (*Gfra1*+/*Kit*-, Cluster II and III). This analysis revealed down-regulation of SSC self-renewal related genes, such as *Etv5* and *Ccnd2*, as well as embryonic stem (ES) cell self-renewal regulator *Tcl1* (Figure 6A) (Table S4). GSEA analysis showed a strong enrichment of glial cell line-derived neurotrophic factor (GDNF) regulated genes in *Gfra1*+/*Kit*- cells, which confirmed the loss of responsiveness to GDNF regulation in *Gfra1*-/*Kit*- cells (Figure 6B) (Oatley et al., 2006). On the other hand, expression of differentiation regulator *Rarg* was increased in *Gfra1*-/*Kit*- cells, which might contribute to the heterogeneous differentiation competence in response to retinoic acid (RA) that resembled steady-state spermatogenesis in adult (Ikami et al., 2015). We found genes related to Ras signaling pathway (FDR= 1.088E-3) enriched in *Gfra1*+/*Kit*- cells, which is consistent with previous finding (Lee et al., 2009). Moreover, a significant enrichment for cell-cell communication and GnHR, as well as neurotrophic signaling related genes were also observed (Figure S8). Functional annotation of genes upregulated in *Gfra1*-/*Kit*- revealed enrichment for DNA methylation e.g. *Tdrd5, Dnmt3b*, and *Ehmt2* (*G9a*), consistent with its role in spermatogonia differentiation (Figure 6A) (Shirakawa et al., 2013). From the pseudotime analysis, it is evident that the transition from stem cells and progenitors to differentiation-primed cells entails a greater magnitude of the changes in gene expression (e.g. complete loss of *Gfra1* and *Etv5* expression). GO analysis indicated other factors involved in mTOR pathway (*Eif4ebp1*, *Pdcd4*, *Cycs*, and *Prkab2*), which is in line with enhanced mTOR pathway activity in adult spermatogonia differentiation (Figure 4C) (Hobbs et al., 2010). Notably, *Nsbp1* and *Eno1*, which have been implicated in cell differentiation in other systems, were also elevated (Figure 4C) (Diaz-Ramos et al., 2012; Rochman et al., 2010).

**Figure 6.**
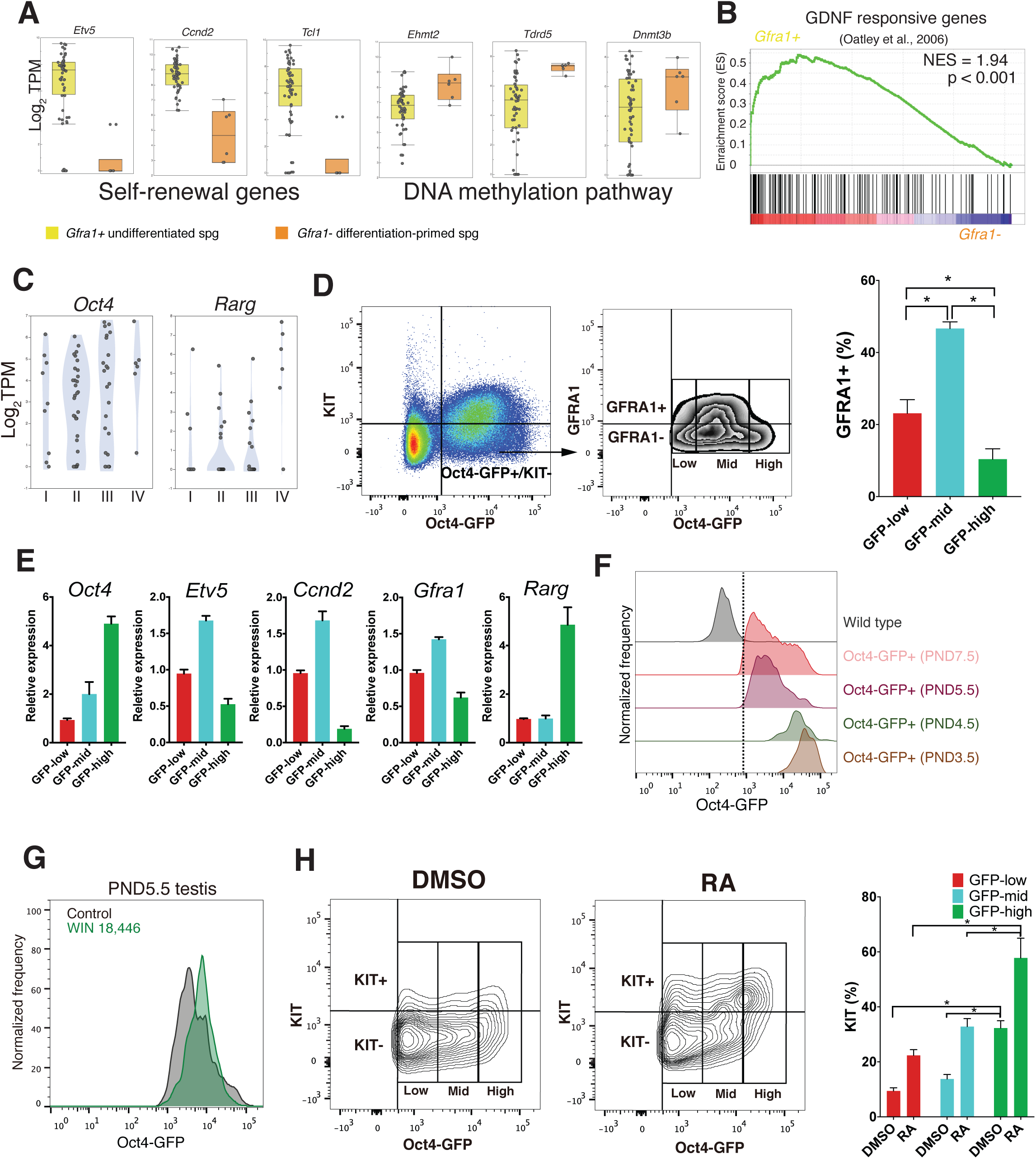
Differentiation-primed cells display enhanced RA responsiveness. (A) Box plots for representative genes identified as differentially expressed (FDR<0.05) between *Gfra1*+ (Cluster II and III) and *Gfra1*-(Cluster IV) spermatogonia. (B) Enrichment plots from GSEA analysis using GDNF-responsive gene sets. Note the significant upregulation of GDNF-responsive genes in *Gfra1*+ cells. (C) Both *Oct4* and *Rarg* mRNA were higher expressed in Cluster IV. (D) FACS sorting scheme to isolate subpopulations of germ cells expressing various levels of Oct4-GFP. (E) Patterns of gene expression measured by RT-qPCR in subpopulations of Oct4-GFP+/KIT- cells sorted by level of Oct4-GFP fluorescence. The population with high Oct4-GFP displayed significantly higher expression of *Rarg*. (F) The Oct4-GFP expression is dynamic throughout the spermatogenesis initiation. Oct4-GFP cells were isolated 3.5-7.5 days after birth and analyzed by FACS. The dashed line represents the cutoff for the determination of Oct4-GFP+ v.s. Oct4-GFP-based on wild-type animal controls. (G) The effect of retinoid signalling on Oct4-GFP expression *in vivo*. Histograms of measured intensity of Oct4-GFP in WIN18,466-treated and control testis. KIT+ cells, Oct4-GFP-low, and Oct4-GFP-high cells were significantly reduced upon WIN 18,446 treatment. (H) Significantly enhanced RA response in Oct4-GFP-high cells. Left panel, the representative plot showing KIT staining in GS culture derived from Oct4-GFP+/KIT- cells. Spontaneously differentiating KIT+ cells mainly reside in Oct4-GFP-high subpopulation. Middle panel, the representative plot showing KIT staining in GS culture after RA-induced differentiation. RA treatment induced KIT expression, which is more profound in the Oct4-GFP-high subpopulation. Right panel, quatification of percentage of KIT+ cells in each group.

### Elevated *Oct4* expression signifies early spermatogonia differentiation and enhanced RA responsiveness

O*ct4* has previously been shown to be crucial for SSC maintenance because *Oct4* is down-regulated by RA-induced differentiation and the knockdown of *Oct4* results in differentiation (Dann et al., 2008). A priori, it might be expected that *Oct4* expression level would decrease along the differentiation process. Surprisingly, *Oct4* was higher expressed in differentiation-primed cells compared to the other subsets (Figure 6C). Given that indexed FACS sorting showed that GFP intensity from the Oct4-GFP transgene positively correlates with endogenous *Oct4* expression at single cell level (Figure S4B and S4D), we next explored whether we could experimentally isolate and characterize the differentiation-primed cells using the level of Oct4-GFP intensity (Figure 6D). Consistent with the prediction from single cell analysis, germ cells sorted by FACS with higher levels of GFP fluorescence had significantly higher expression of *Oct4* and *Rarg* (Figure 6E). To further associate Oct4-GFP expression to GFRA1 protein level, we pregated on Oct4-GFP+ cells and then examined GFRA1 staining in different subsets and found that GFP-high gated cells contained significantly fewer GFRA1+ cells (Figure 6D).

As an independent means to confirm that Oct4-GFP-high cells are primed to differentiation *in vivo*, we traced Oct4-GFP expression through developmental stages, reasoning that the earliest differentiation-primed population would expand during development. We observed that gonocytes expressing strong Oct4-GFP cells at PND3.5 and PND4.5 largely disappear by PND5.5, while another cell population with relatively lower Oct4-GFP level emerged at PND5.5, which represent newly formed spermatogonia. Importantly, PND5.5 germ cells exhibiting higher GFP level expanded in number and this subset became more apparent at PND7.5 (Figure 6F). We reasoned that blockage of differentiation of spermatogonia after PND4.5 should keep them in an undifferentiated state and lead to the reduction in the proportion of cells primed to differentiation but not affect early gonocyte development. We treated PND4.5 Oct4-GFP mice with WIN 18,446, which chemically inhibits RA synthesis and retains spermatogonia at their SSC state (Agrimson et al., 2017; Paik et al., 2014). As expected, KIT expression was down-regulated upon treatment, suggesting a lower level of RA activity and blockage of differentiation (Figure S9). Indeed, WIN 18,446 treated PND5.5 testis are essentially devoid of cells expressing the highest Oct4-GFP present in normal PND5.5 and PND7.5 testis (Figure 6G). In addition, Oct4-GFP-low cells expressing minimal level of GFRA is also decreased after WIN 18,446 treatment, implying that they are going through differentiation.

We next explored whether Oct4-GFP+ cells maintained *in vitro* would recapitulate this differentiation priming process, and whether Oct4-GFP-high cells displayed higher differentiation competence in response to RA. To test this, primary cultures derived from PND5.5 Oct4-GFP+/KIT- cells were treated with all-trans RA or vehicle, followed by examination of KIT expressin across cell subsets with different Oct4-GFP intensity. Control cultures without RA exposure showed signs of spontaneous differentiation as a small fraction of cells began to express KIT. Interestingly, KIT+ cells primarily reside in Oct4-GFP-high cells (Figure 6H). RA exposure led to an increased proportion of cells positive for KIT and the percentage of Oct4-GFP-high cells that became KIT+ was significantly higher when compared to Oct4-GFP-low and Oct4-GFP-mid cells (Figure 6H).

Collectively, these data show that Oct4-GFP-high cells display molecular and functional properties as cells primed to differentiation both *in vivo* and *in vitro*. Elevation of *Oct4* expression thus reflects a switch from self-renewing to differentiation commitment.

## DISCUSSIONS

Previous single-cell gene expression analysis on neonatal spermatogonia at PND6 is restricted to only 172 genes, which limits the resolution to dissect the molecular heterogeneity (Hermann et al., 2015). In this study, we extended this analysis to whole-transcriptome profiling on Oct4-GFP+/KIT- cells from PND5.5 testes, which represent a broader undifferentiated germ cell population. In agreement with the recent study (Song et al., 2016), this approach allowed us to reveal early germ cell development trajectory in which migratory gonocytes transition into undifferentiated spermatogonia that subsequently proceed to differentiation.

Importantly, our study provides the global view of gene expression dynamics underlying GST, which permits elucidating important events that dictate the fate of gonocytes. First, we discovered several molecules that might be potentially involved in gonocyte migration. Second, GST is accompanied by a global activation of transcription and change of cell cycle status. Third, our results showed that apoptosis-related genes are actively regulated during gonocyte transition. Particularly, transitional gonocytes seem to have a lower threshold for initiating apoptosis (elevated death receptor pathway) and increased susceptibility to DNA damage-induced apoptosis (reduced DNA repair pathway). This raises the possibility that transitional gonocytes possess the poised apoptotic machinery, which renders them more vulnerable to apoptotic cell death. Further studies using our resources will help shed light on the mechanistic nature of how signals regulate gonocyte cell cycle and migration during GST.

Globally, our data explicitly pinpointed many developmental hallmarks in the literature previously assumed to exist in neonates based on studies in adult spermatogenesis (Figure S11A). In corroboration of a recent ‘hierarchinal model’ of adult SSC, we showed that neonatal primitive stem cell and progenitor populations are distinguished by a gradient expression of SSC maintenance genes (*Id4, Gfra1, Etv5* and *Tspan*8) and differentiaoin-promoting genes (*Sohlh1* and *Rarg*) (de Rooij, 2017), which closely reminiscent with the recent single-cell analysis in human adult SSCs (Guo et al., 2017)). The ‘ultimate’ stem cells proposed in this model might reside in the primitive stem cell population we identified and would be enriched by cell surface markers that are specific to this population. Among the panel of cell-surface markers we described, the expression of *CD87* is limited to the neonatal stage. It is tempting to speculate that *CD87* may play a pivotal role in the intial establishment of SSC pool, potentially through the regulation of migration.

Our results have implications for understanding how neonatal SSCs respond to RA and differentiate in vivo. A recent study demonstrated that adult SSCs that are labelled by ID4-GFP+ or GFRA1+ express Rarg. Consistent with this finding, we observed Rarg expression in both Cluster I and II cells, albeit at lower level. We further showed the transition from Gfra1+/Rarg-low (Cluster II and III) to *Gfra*1-/*Rarg*-high (cluster IV) cells signifies the early spermatogonia differentiation. Higher *Rarg* expression may enhance the sensitivity of these cells to RA exposure, which closely resembles the recently described differentiation model in adult spermatogenesis (Ikami et al., 2015). This subset of differentiation-primed cells notably displays a surprisingly higher level of Oct4. It has been shown that Oct4 positively controls the level of RARγ and directly regulates retinoic acid pathway in ESC to direct the differentiation process (Niwa et al., 2000). This raised the possibility that it is also crucial for SSCs to maintain Oct4 at an appropriate level in order to preserve self-renewal capacity, and change in Oct4 level is able to specify self-renewal and differentiation cell autonomously. The detailed mechnisam underlying this observation awaits for further invesitigation. Nevertheless, our finding suggests Oct4-GFP can be used as an excellent reporter system for real-time visualization of changes in cellular state during SSC differentiation.

At the same time, this high-resolution data set provides a more comprehensive characterization of molecular cascades underlying early spermatogonia differentiation (Figure S11B). Our analysis showed that the regulation of energy metabolism is tightly coupled to spermatogonia differentiation. Genes related to lipid metabolism are highly enriched in stem cell pool, which extends previous findings on the role of fatty acid metabolism in stem cell maintenance (Ito and Suda, 2014). Our results also indicated that the ribosome biogenesis increased in progenitors compared to stem cell spermatogonia. As translation rate is correlated positively with ribosome biogenesis, our results implied an enhanced need for global protein synthesis during differentiation. Interestingly, increased protein synthesis has been observed in GSC development in *Drosophila* (Sanchez et al., 2016). Investigating the cell cycle heterogeneity further demonstrated that differentiation of spermatogonia is intertwined with cell cycle states. Interestingly, cell cycle regulator *Ccnd2* was identified as the hub gene in self-renewal module (Figure S6C), which strongly suggested that it might serve as an important modulator for the coupling of spermatogonia differentiation and cell cycle progression. Our results, therefore, strengthened the hypothesis that faster cell cycle progression and a shorter G1 phase in stem cell pool could be accomplished by induction of cyclin D expression as reminiscence of the interconnection of cell fate decision and cell cycle demonstrated in other stem cell systems (Becker et al., 2006).

This study also provided an opportunity to improve our understanding of developmental disease processes, further attesting the importance of our work. Aberrant expression of *Ccnd2* is believed to contribute directly to the formation of testicular germ cell tumors (TGCTs) (Rodriguez et al., 2003). Similarly, another hub gene *Spry4* has been proposed as one of the susceptible genes for TGCTs (Karlsson et al., 2013). Moreover, the expression of CD87 was significantly increased in TGCT, which may be relevant in testicular cancer progression (Ulisse et al., 2010). On the other hand, low level of ERCC1 or recruitment of FADD is likely to be key determinants of cisplatin sensitivity in TGCT (Spierings et al., 2003; Wood, 2011). Since defective gonocyte differentiation might be the origin of testicular germ cell tumors, it is intriguing to speculate that intrinsic gene expression phenotype (Ercc1^low^ and Fadd^high^) of the transitional gonocytes may be retained in TGCTs and plays a significant role in determining the cisplatin sensitivity phenotype.

In conclusion, we envisaged that our comprehensive data set and strategy would serve as a powerful tool to further dissect the heterogeneity in adult human and mouse spermatogonia, paving the way for identification of molecular mechanisms underlying adult male germ cell development and associated diseases in an unbiased and precise manner.

## MATERIALS AND METHODS

### Immunofluorescence staining

Immunofluorescence staining of testes sections were performed as described previously (Oatley et al., 2011) in conjunction with the antibodies against KIT, GFRA1, and PLZF. Images were acquired using Leica SP2 confocal microscope.

### RNA sequencing

Undifferentiated germ cells from PND5.5 testes were purified using the method described previously (Garcia and Hofmann, 2012) in conjunction with KIT antibody. Isolated cell were captured using the Fluidigm C1 Auto Prep system. cDNAs were prepared using the SMARTer Ultra Low RNA Kit for Illumina (Clontech). Libraries were generated using the Nextera XT (Illumina) protocol. RNA-Seq libraries were sequenced using Illumina HiSeq 2000. Data is available in the public repository GEO (GSE107711).

### Analysis of RNA-Seq data

Raw reads were pre-processed with trim galore for trimming Illumina adaptor sequences and mapped to mouse mm9 genome. We estimated expression level for all RefSeq genes using Salmon (Patro et al., 2017) and selected the 500 top-ranked PCA genes for clustering and PCA analysis. Pseudotime ordering were performed with Monocle (v1.2.0) as previously described (Trapnell et al., 2014a). Differential gene expression analysis was performed with the Bayesian approach to single-cell differential expression (SCDE) (Kharchenko et al., 2014) or DESeq2 method (Love et al., 2014). R package “WGCNA” was used to construct the weighted gene co-expression network. GO enrichment was performed using DAVID (Huang da et al., 2009), ToppGene Suite (Chen et al., 2009) or ClueGO (Bindea et al., 2009).

## AUTHOR CONTRIBUTIONS

JL and TL designed the study, performed experiments, analyzed and interpreted data, and prepared the manuscript. SN participated in experimental design, single-cell capture experiment and manuscript writing. JT, AL, and YQ provided intellectual inputs. NT, WC and PF advised on data interpretation. All authors read and approved the final manuscript.

## ACKNOWLEDGEMENT

The authors acknowledge the support of Core laboratory in School of Biomedical Sciences, The Chinese University of Hong Kong and Prof. Bo Feng (School of Biomedical Sciences, Faculty of Medicine, The Chinese University of Hong Kong) for providing Oct4-GFP transgenic mice. This work was supported by General Research Funds from Hong Kong Research Grant Council (Project no. 468312, 469713) and Lo Kwee-Seong Biomedical Research Seed Fund from School of Biomedical Sciences, The Chinese University of Hong Kong (Project no. 7104687).

